# Studies on squalene biosynthesis and the standardization of its extraction methodology from *Saccharomyces cerevisiae*

**DOI:** 10.1101/309609

**Authors:** Kalaivani Paramasivan, Kavya Rajagopal, Sarma Mutturi

## Abstract

**Aims:** The present work focuses on studies on squalene improvement in the *S. cerevisiae* and development of squalene extraction procedure based on mechanical disruption of cells.

**Methods and Results:** In this study, a homogenization-based extraction method was developed and was compared to five conventional methods of squalene extraction. Squalene recovered from this novel procedure gave 3.5– fold, 10-fold, 16-fold and 8.1-fold higher yield than standard procedures viz., saponification with 60% KOH, acidic saponification, saponification with 18% KOH and glass beads method, respectively. Furthermore, this procedure has been evaluated on laboratory *S. cerevisiae* strains such as BY4742 and CEN.PK2-1C (native), deletion strains (*ERG6* and *ERG11*) and *tHMG1* overexpressed *S. cerevisiae* strains. When sonication method of cell lysis was replaced with homogenization it was found that the yields were significantly higher and reached a value of 9 mg/g DCW in case of BY4742. In addition, Squalene yield in ergosterol mutant strains has been analyzed and was found to be 1.8-fold and 3.4-fold higher in *ERG6* and *ERG11* deletion strains, respectively, than BY4742. Squalene was also found to be higher at the optimized temperature of 30°C and pH 6.0. Furthermore, tolerance of *S. cerevisiae* to external squalene at various concentration has been carried and found that the organism was tolerant up to 25 g/L of squalene.

**Conclusions:** Homogenization based mechanical disruption was observed to yield higher squalene and the SEM analysis corroborates these findings. The synergistic effect of *ERG11* downregulation and *tHMG1* over-expression has led to significant increase in squalene yield.

**Significance and Impact of the Study:** Sonication and homogenization has been used as a cell-disruption method for the first time in squalene extraction and squalene yield from homogenization method of cell lysis in BY4742 has been found to be 9 mg/g DCW which is the highest reported so far in a wild-type strain.

## Introduction

Squalene is one among the diversified terpene molecules synthesized in yeast cells (Paramasivan and Mutturi 2017a). Squalene is industrially hydrated to squalane which is used as an ointment as well as a lubricant in automobiles. Squalene can be used as a drug carrier and as an adjuvant in vaccines (Naziri et al. 2011). It is used as a moisturizer or emollient in cosmetic industry and it has been proven to decrease UV-induced DNA damage (Kohno et al. 1995). Squalene is involved in tumor-suppression (Rao et al. 1998) as well as in immunity enhancement (Kelly 1999). It exhibits antibacterial, antifungal (Smith 2000) and antivirulent activity (Sri Charan Bindu et al. 2015). In addition, it also has cholesterol-lowering effects and used in the treatment of cardiovascular diseases (Dhandapani et al. 2007). Hence it has been widely used as a dietary supplement. Further, owing to the advantage of having a hydrocarbon backbone, it has proven itself as a potential source of biofuel (Bergquist et al. 2014). Squalene is less accumulated in yeast strains owing to the tight-regulation in the ergosterol biosynthesis pathway and the storage inability of lipid droplets (Polakowski et al. 1998). Liver oil of sharks (*Squalus spp.*) is the richest source of squalene with its composition reaching upto 40-70% of its dry weight (Gershbein and Singh 1969). The plant oil sources include amaranthus seed oil, wheat germ oil, rice bran oil and olive oil (Spanova and Daum 2011). However, shark liver oil is not a sustainable source for squalene due to imminent danger of extinction and also because of prevailing concern over the leaching of pollutants. Microbial squalene hyper-accumulaters *Pseudozyma sp.* and *Aurantiochytrium sp.* accumulates upto 7% and 32% of their dry cell weight (Katabami et al. 2015). The natural squalene accumulating microbes has a longer cultivation cycle, low squalene yield and productivity, and does not have well established gene manipulation techniques and growth cultivation conditions in industrial scale. *S. cerevisiae* is one of the most sought out organism for squalene synthesis as it accumulates around 1 to 4.6% per dry cell weight of ergosterol depending on the strains (Arnezeder and Hampel 1990). Squalene being a precursor molecule in ergosterol pathway is also produced in higher amounts by the yeast strains. Genetic manipulation techniques have been applied in *S. cerevisiae* for improving the flux towards mevalonate pathway which in turn increases isoprenoid synthesis.

Kamimura *et al.* (1994) has pioneered the gene disruption in ergosterol biosynthesis pathway for squalene accumulation and has reached a squalene yield of 5 mg/g dry cell weight by random mutation in *ERG1* gene. Bhattacharjee *et al.* (2001) has produced squalene in a commercial *S. cerevisiae* strain with the yield of 41 μg/g DCW under anaerobic fermentative conditions. Garaiová *et al.* (2014) has studied the squalene enhancement in *ERG1* mutant strains in the presence of ergosterol biosynthesis inhibitor, terbinafine and has achieved a squalene yield of 1000 μg per 10^9^ cells. Mantzouridou, Naziri and Tsimidou (2009) has achieved a maximum squalene yield of 3.1 mg/L under optimized semi-anaerobic, inoculation and bioprocess culture conditions. Over-expression of truncated version of HMG-CoA reductase is one of the most widely used strategy for squalene accumulation and for improving the flux towards mevalonate pathway in *S. cerevisiae.* Hmg1p undergoes feedback-inhibition by the end product ergosterol. To overcome this, a truncated version of the enzyme has been over-expressed and tHmg1p overexpression has enhanced squalene improvement (Donald et al. 1997; Polakowski et al. 1998).

The extraction of squalene is critical as it is sequestered in different organelles including lipid droplets, membranes and endoplasmic reticulum and there is a need for its efficient recovery from the cell. Squalene has been conventionally extracted employing pyrogallol and saponification methods using potassium hydroxide (Tokuhiro et al. 2009; Mantzouridou and Tsimidou 2010; Shin et al. 2012; Hull et al. 2014). Estimation of squalene is very critical as it is produced intracellularly and inefficient extraction of squalene from the *S. cerevisiae* cells might lead to erroneous selection of sub-optimal strains. The genetic modification increases the capacity of squalene production intracellularly and a robust analytical procedure is required to quantify this metabolic capacity of *S. cerevisiae* strain.

Yeast cell walls are known to have a rigid cell-wall and requires harsh treatments for the breakage of cells. The cell lysis could be achieved through enzymatic methods such as lysozymes and proteinases treatment, chemical treatments such as sodium dodecyl-sulphate, EDTA, phenol and guanidium thiocyanate; heat treatments such as autoclaving and microwaving; mechanical methods such as sonication and homogenization. Freeze-thaw cycles and freeze-boil cycles has also been efficient in cell disruption. Mechanical methods are easy to scale-up for large-scale cultivations, relatively cheaper and could be applied at industrial level. Ultra-sonication and high-pressure homogenization are some of the most commonly used cell-disruption methods for intracellular product extraction. Here in the present study, mechanical-based methods such as sonication and homogenization of yeast cells were developed for efficient recovery of squalene and was compared with traditional saponification techniques. All the selected analytical procedures were enumerated using *S. cerevisiae* laboratory strains with and without *tHMG1* overexpression.

## Materials and methods

### Media and culture conditions

Euroscarf yeast strains BY4742, CEN.PK2-1C, *ERG*11*Δ*, *ERG*6Δ and *E.coli* strain DH5α were kindly provided by Prof. Ram Rajasekharan (Lipid Science Dept., CSIR-CFTRI). Squalene standard was procured from Sigma-Aldrich (Bangalore, India). The restriction enzymes, ligase and phusion DNA polymerases were obtained from New England Biolabs, (Ipswich, MA, USA). Yeast strains BY4742, CEN.PK2-1C, *ERG11Δ*, *ERG6Δ* and recombinant strains were grown at 30°C either in YPD or synthetic medium without uracil(Paramasivan and Mutturi 2017b). *E. coli* DH5α was grown at 37°C in LB Media (10 g tryptone/L, 10 g yeast extract/L and 5 g sodium chloride/L) in the presence or absence of ampicillin (100 μg/mL) as per requirement and is used for the propagation of plasmids. Cell growth was determined by measuring the OD_600_.

### Plasmids and strain construction

The expression plasmid used in the study is pYES2-NT-C (URA3,2μ,P_Gal1_); pYGH developed from pYES2-NT-C as reported in previous study(Paramasivan and Mutturi 2017b) was introduced into the parent strains BY4742, CEN.PK2-1C, *ERG6*Δ and *ERG11*Δ to obtain SK2, SK3, SK4 and SK5 strains (Table 1), respectively, using Sc easy Com transformation kit (Invitrogen, Bangalore, India). All DNA sub cloning steps were performed with *Escherichia coli* DH5α using standard bacterial transformation methods as described by Sambrook and Russell (2001)^29^ and sequences of cloned genes were verified by sequencing. Yeast transformants were picked and pre-cultured in 50 mL of yeast synthetic culture at 30°C, 180 rpm for 16 h.

**Table 1:**
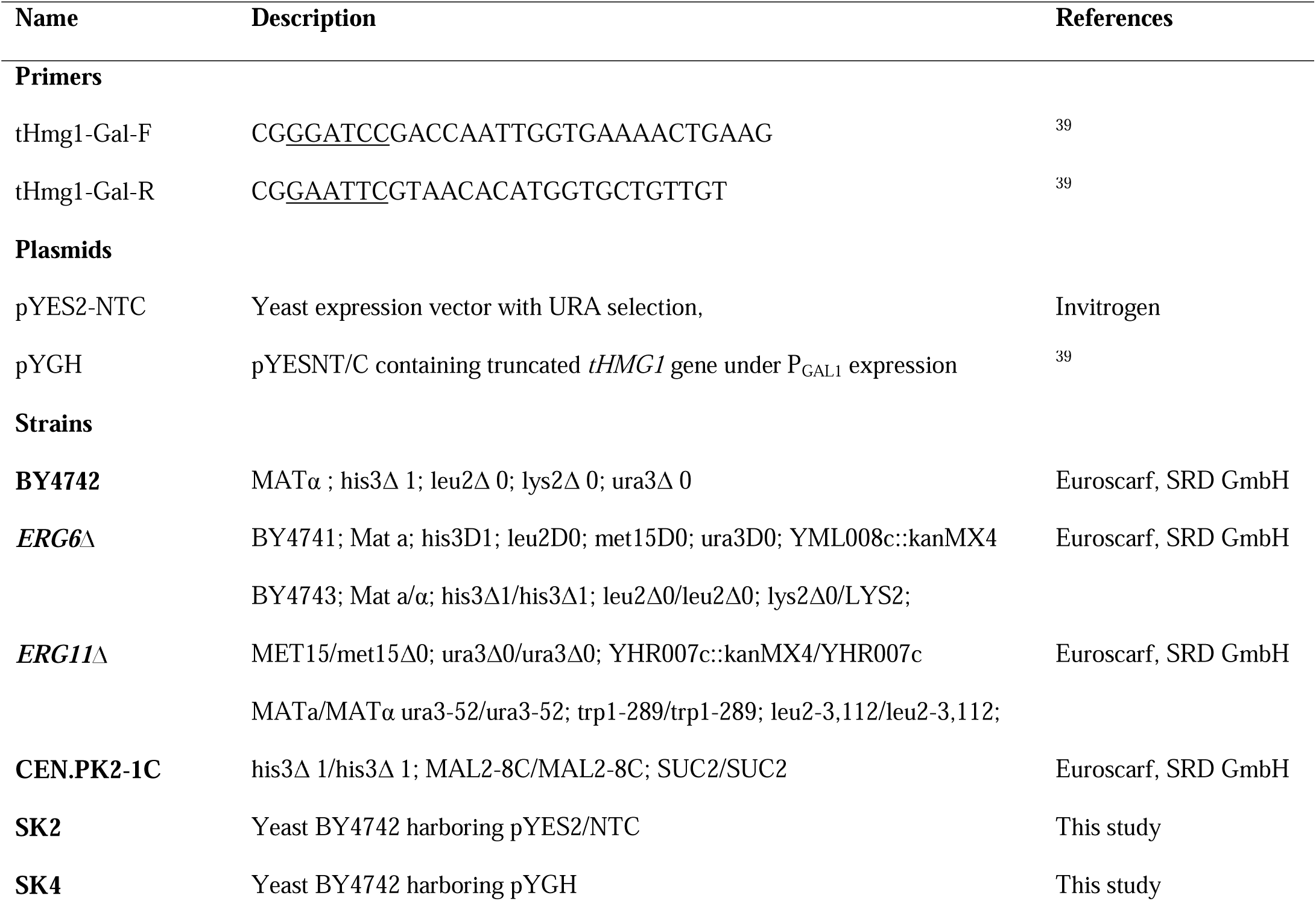
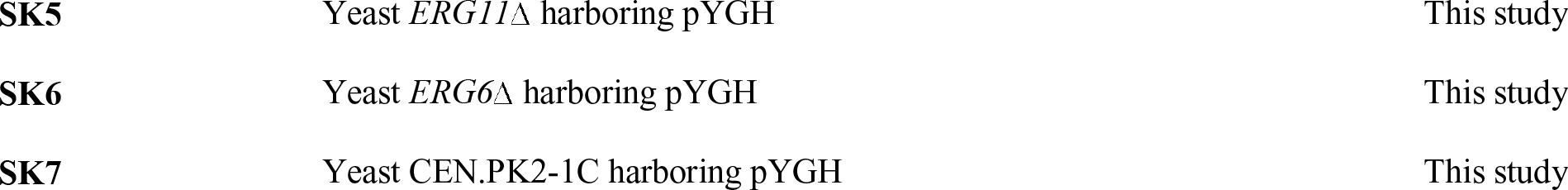
Primers, plasmids and strains used in the study

### Squalene extraction methods and quantification

#### Extraction method for squalene by mechanical disruption

Yeast cells were grown in pre-culture yeast synthetic medium for 15 hours. The inoculum from pre-culture was used in production culture medium at an initial OD of 0.1 and were grown for 24 hours at 30°C, 180 rpm. The cells were harvested at 5684 g for 10 minutes at 4°C and later frozen at -20°C. The cells were subjected to lyophilization after deep-freezing for 2 hours. The samples were freeze-dried at -50°C under vacuum for 4 hours. The freeze dried cells were then dispersed in 30 mL of chloroform : methanol solution (2:1, v/v) and were disrupted either by sonication/homogenization. Homogenization was carried out using SCILOGEX D-160 homogenizer (Rocky Hill, Connecticut, USA) for 5 minutes under ice-cold conditions. The cell lysates were then subjected to overnight extraction at 30°C, 180 rpm in shake flasks while the volume of organic solvent mixture of chloroform : methanol was made up to 50 mL. The samples were filtered using Whatman’s No.1 chromatography filter paper after extraction and was subjected to flash evaporation. Flash evaporation of solvents was carried out under vacuum in round bottomed flasks at 50°C and 300 psi pressure. The samples were then dispersed in 500 μl of chloroform and stored at -20°C. The samples were then subjected to high speed centrifugation to remove debris prior to RP-HPLC analysis. Sonication-based extraction has been carried according to the protocol described in Paramasivan and Mutturi (Paramasivan and Mutturi 2017b).

#### Saponification and glass-bead method of extraction

##### Extraction by saponification with 60% KOH at 45°C(Mantzouridou and Tsimidou 2010) (Method-1)

Yeast cell pellets were suspended in 5 mL of 60% KOH, 7.5 mL of methanol and 7.5 mL of pyrogallol (0.5% w/v). The mixture was incubated overnight in a shaker incubator at 45°C. The solution was then subjected to extraction using 10 mL of hexane by centrifugaton at 5684 g for 10 minutes. The upper layer was collected and vacuum dried at 40°C in a vacuum oven until the solvent is completely evaporated. After the completion of vacuum drying the tube was added with 1 mL of chloroform to resuspend the fraction.

##### Extraction by acidic saponification at 70°C (Hull et al. 2014) (Method-2)

Cell Pellets were suspended in 2 mL of 0.5 N HCl and treated for 20 minutes at 100°C and later cooled to room temperature. 3 g of KOH, 12.5 mL of methanol with 0.025 g of pyrogallol in it was later added to cooled suspension and incubated at 70°C for 2 hours. The contents were subjected to extraction using 10 mL of Hexane and vacuum dried at 40°C until complete solvent evaporation and the samples were resuspended in 1 mL of chloroform.

##### Extraction by saponification with 18% KOH at 90°C (Shin et al. 2012) (Method-3)

Cell pellets were suspended in 10 mL of 7:3 methanol: water with 18% KOH, 0.1% Pyrogallol. The contents were then heated at 90°C for 2 hours. Further, the contents were subjected to extraction using 10 mL of hexane followed by vacuum drying at 40°C and re-supended in chloroform.

##### Extraction by glassbeads method using pentane (Tokuhiro et al. 2009)

The yeast suspension of 2 mL was pelleted down and dispersed in 0.4 g of glass beads containing 0.2 mL of TE buffer and later subjected to vortexing at room temperature for 30 minutes. Initially methanol and pentane (1:5) extraction was carried followed by pentane only (two times) and the organic layer was collected and dried. The extracts were suspended in chloroform prior to HPLC analysis.

### Analytical procedures

RP-HPLC was performed isocratically using solvent delivery system consisting of two HPLC pumps and a rheodyne injection valve with a 20 μl of fixed loop (Shimadzu Scientific instruments, Kyoto, Japan). The chromatograms were processed using LC Solution software. Squalene analysis was carried using a semi preparative reversed-phase C-18 column (Phenomenox Kinetex, Hyderabad, India) (particle size 5 μm, 250 X 4.6 mm i.d.) maintained at a temperature of 35°C. The chromatograph was coupled to a UV-VIS Diode array multiple wavelength detector. The flow rate was set at 1.5 mL/min. The mobile phase used was 100% acetonitrile. The injection volume was set at 20 μl. Detection and quantification of squalene was carried out at 195 nm. Peak identification was achieved by comparing the retention time with the authentic standards and confirmed by spectral data. Quantification was accomplished with the help of standard curves calculated using linear regression analysis. To obtain the dry cell weight (DCW), cells harvested by centrifugation from the 50 mL culture were oven-dried and measured. All optical densities at 600nm (OD600) measurements were taken using a Shimadzu UV-2550 spectrophotometer.

### Thin layer chromatography

The lipid extracts containing squalene and ergosterol in chloroform and ethanol, respectively, were subjected to qualitative thin layer chromatography in silica gel using two solvent systems. Commercial plates precoated with silica gel (60G F  ) were procured from Merck chemicals, (Bangalore, India). Cyclohexane has been used as a solvent system for squalene analysis while petroleum ether: diethyl ether: acetic acid in the ratio 90:10:1 has been used as a solvent system for ergosterol analysis. Once the mobile front reached the desired height on the silica, the plates were removed from glass chamber and were exposed to 20% H_2_SO_4_ mist and later incubated for heating at 70°C in hot air oven for visualization. The bands were also visualized in UV chamber as the squalene has an absorbance of 195 nm. The spot intensity was compared with the squalene standard obtained from Sigma-Aldrich (Bangalore, India).

### Scanning electron microscope

The cell lysis was visualized under scanning electron microscope. The chemical fixation of cells were carried out using 2.5% glutaraldehyde. The cells were pelleted down and washed with 0.1 M phosphate buffer (pH 7.2). The washed cells were treated with 2.5% of glutaraldehyde and incubated at 4°C overnight. The fixed cells were pelleted down and washed serially with 10%, 20%, 50%, 70% and 100% ethanol. The samples were kept in desiccator for further analysis. Following dehydration, critical point drying was carried out. The samples were sputter coated with gold before subjecting to scanning electron microscopy. Finally, SEM micrographs of critical-point dried cells were generated using Leo Scanning Electron Microscope Model 435VP (Leo Electron Microscopy Ltd., Cambridge, UK).

## Results

### Effect of mechanical disruption of cells on squalene extraction

Squalene being an intracellular compound, cell lysis becomes an imperative step for its extraction. Initially four different laboratory strains (BY4742, CEN.PK2-1C, *ERG11*Δ and *ERG6*Δ) were used for comparing the chemical (saponification, 60% KOH at 45^°^C) and mechanical (sonication)-based extraction procedures for squalene from the cell pellet. The deletion strains *ERG11Δ* and *ERG6Δ* were selected based on the rationale that the deletions downstream of squalene towards ergosterol could improve intracellular squalene accumulation. From Figure 1A it can be observed that the squalene titer varied across method-used for extraction and also the *S. cerevisiae* strain used for its synthesis. It was found that sonication-based squalene extraction yielded higher titers than saponification across all the tested strains. It can also be observed that the squalene produced using CEN.PK2-1C is higher than BY4742 strain. It was suggested that CEN.PK strains accumulate higher ergosterol in comparison to BY series(Daum et al. 1999). Similarly based on the rationale for selecting mutant strains, it has been validated that *ERG11Δ* and *ERG6Δ* accumulate higher squalene in comparison to parent BY series. From this experiment it is evident that sonication-based squalene extraction is efficient over saponification. In order to understand the possibility of having higher squalene extraction based on mechanical-method, two more methods viz., glass beads - and homogenization-based cell lysis were investigated on BY4742 strain. As there were several modifications to standard saponification method for extracting squalene in the existing literature, three such variants were included in the experiment to understand variations amongst these methodologies in terms of squalene extraction.

**Figure 1.**
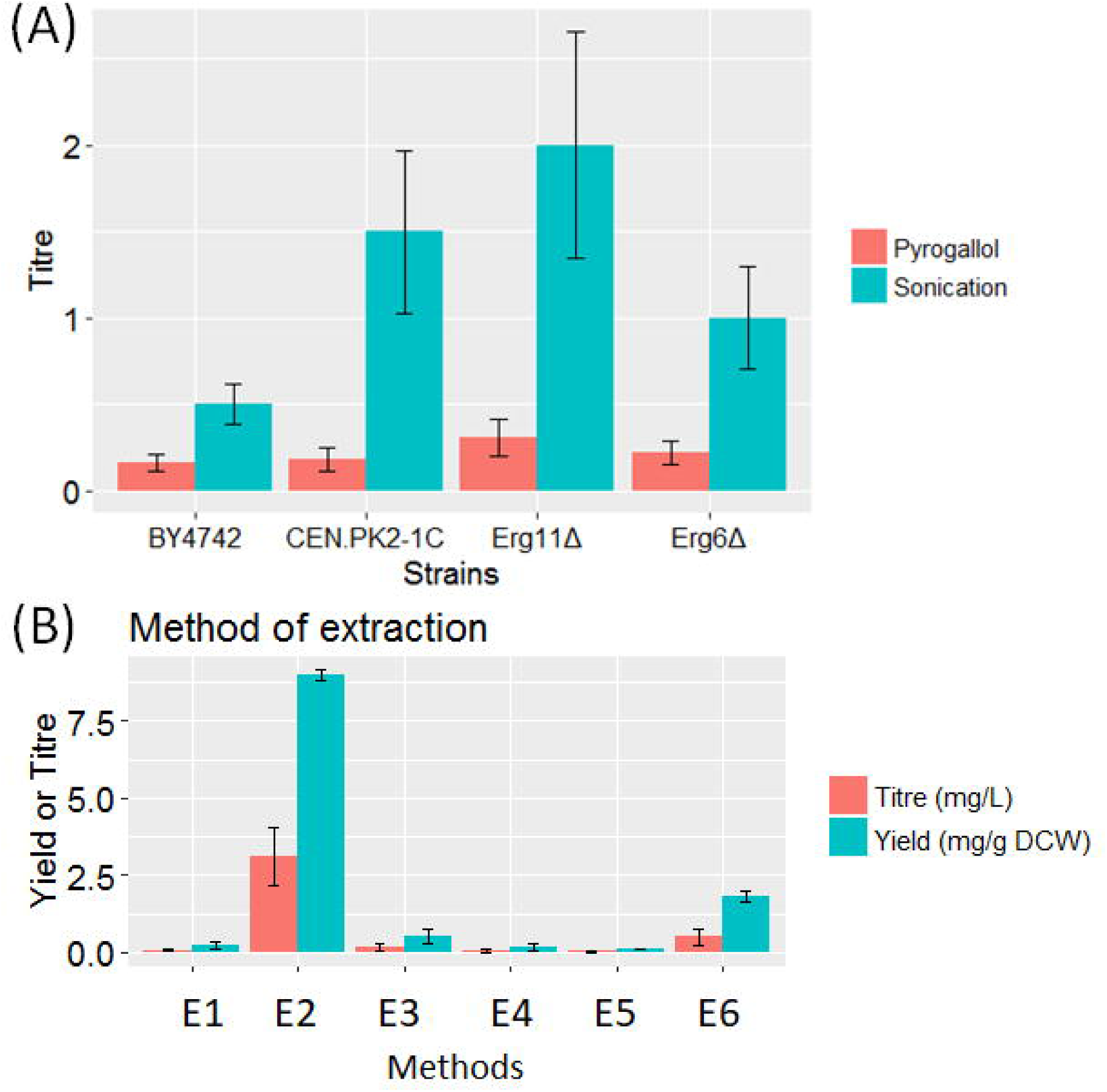
(A). Comparison of squalene yield from various strains using sonication and saponification with 60% KOH at 45°C. (B). Squalene yield in BY4742 using different methods of extraction.

Among these four different methods, homogenization-based extraction has significantly improved squalene extraction than chemical saponification and glass beads methods (Figure 1B). Also it can be observed that saponification method-1 fared better than other two saponification methods. The devised method has shown 3.5-fold, 10-fold, 16-fold and 8-fold higher than the methods reported with Pyrogallol at 45°C (Mantzouridou and Tsimidou 2010), Pyrogallol at 70°C (Hull et al. 2014), Pyrogallol at 90°C (Shin et al. 2012) and Glass beads method (Tokuhiro et al. 2009) respectively (Figure 1A) (Table 2).

**Table 2:** Efficiency of sonication and homogenization over the standard extraction methods

| Strain | Method of extraction | Yield | Yield fold | Yield fold |
| --- | --- | --- | --- | --- |
|  |  | (mg/g<br>DCW) | increase using<br>sonication | increase using<br>homogenization |
| BY4742 | Sonication | 1.8 | NA | 5 |
|  | Homogenization | 9 | NA | NA |
|  | Extraction by saponification<br>with 60% KOH at 45°C | 0.51 | 3.5 | 17 |
|  | Extraction by acidic<br>saponification at 70°C | 0.18 | 10 | 50 |
|  | Extraction by saponification<br>with 18% KOH at 90°C | 0.11 | 16 | 82 |
|  | Extraction by glassbeads<br>method using pentane | 0.22 | 8.1 | 41 |

617 NA: Not Applicable

Optimal solvent system for extracting squalene was also studied. Experiments using solvents such as hexane, chloroform, methanol, acetone and ethanol ranging from non-polar to polar were conducted. Chloroform was found to be a better solvent for the recovery of non-polar neutral lipid squalene. The squalene extracted using chloroform as a solvent was found to be 2.5 folds higher over the other solvents such as methanol, ethanol, acetone and hexane (Data not shown). Further, the cell lysis by sonication was compared with homogenization and has been observed that homogenization shows higher cell disruption efficiency than sonication. Homogenization method has given a squalene yield of 9 mg/g DCW which is 5-fold higher over the sonication method (Figure 1B). When the homogenization method was compared with standard methods of extraction, the yield was found to be 17.6-fold, 50-fold, 82-fold and 41-fold higher than the standard methods of extraction (Table 2).

Furthermore, the extraction of an intracellular bio-active product relies on the efficient disruption of cells. In the present study, it has been hypothesized that squalene being an intracellular product requires strong mechanical forces for its extraction in the solvents. Hence, we have chosen sonication as a cell-disruption method for squalene extraction. Although squalene is a thermostable compound, when the cell disruption by sonication was carried out at room temperature, the squalene yield was found to be drastically reduced (Data not shown). Hence the sonication has been carried out in cooling water bath under ice-cold conditions. Prolonged sonication has led to squalene degradation rather than the increased release probably due to the high temperature and reduced cooling in the proximity of sonication probe. Horn-type sonicator has been preferred over the bath-type sonicator as the direct immersion of sonicator probe in cell suspensions is relatively more effective in releasing the intracellular products.

The control and the lysed cells were visualized using scanning electron microscope and the micrographs are presented in Figure 2. The morphological variations on the surface of wild-type *S. cerevisiae* cells (Figure 2A) and the lysed cells (Figure 2B and Figure 2C) were observed using scanning electron microscope (SEM). As it can be visualized from the micrographs, the cleaved cells has shown morphological changes than the control cells. Figure 2B and Figure 2C shows a clump of cells lysed by homogenization and sonication, respectively. The wounds on the cell surface are clearly visible in case of homogenized cells (cf. Figure 2C). The microscopic visualization of a single cell broken down by homogenization can also be seen in Figure 2B. From the SEM analysis, it can be concluded that the lysis of cells by homogenization has been found to be more effective than the sonicated cells. SEM micrographs further confirms that homogenization can be used as an efficient method for effective release of intracellular products from yeast cells.

**Figure 2.**
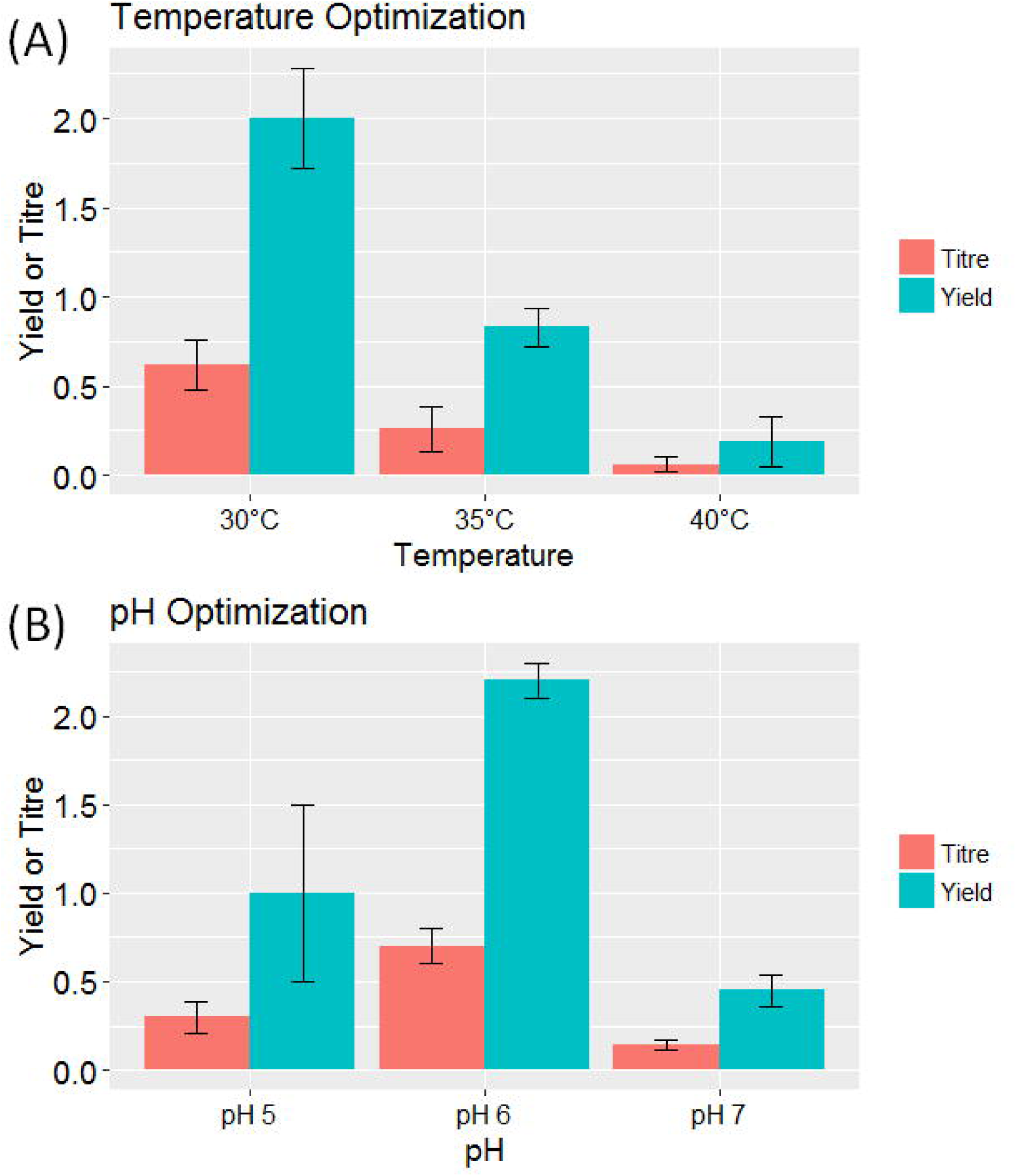
Scanning electron micrographs of *S. cerevisiae* wild-type cells and lysed cells. (A), (B). *S. cerevisiae* wild-type cells, (C). Partially lysed clumps of *S. cerevisiae* cells by homogenization method, (D). Single homogenized cell. (E). Partially lysed *S. cerevisiae* cells by sonication method. (F). Single sonicated cell.

### Effect of pH and temperature in squalene synthesis

Although yeast cells can withstand a wide range of pH and temperature, they achieve their optimum growth at a pH of 5.5 and 30°C. Temperature and pH are the critical parameters for synthesis of a metabolite. To exert a control over the parameters and to achieve the maximum squalene yield, BY4742 wild-type strain was grown at the temperatures of 30, 35 and 40°C and a pH range of 5, 6 and 7 for 24 hours under shaking conditions at 180 rpm. Squalene extraction was carried out at different temperature and pH and it has been observed that the squalene synthesis was higher at the pH of 6 and temperature of 30°C which is in accordance with the published results (Rasool et al., 2016) (Figure 3A). Squalene yield was reduced to its half at 35°C and it was found to be 10 times lower at 40°C. Squalene yield was 2-times higher in pH-6 than pH-5 while it got 4-times lowered in the cells grown at pH-7 (Figure 3B).

**Figure 3.**
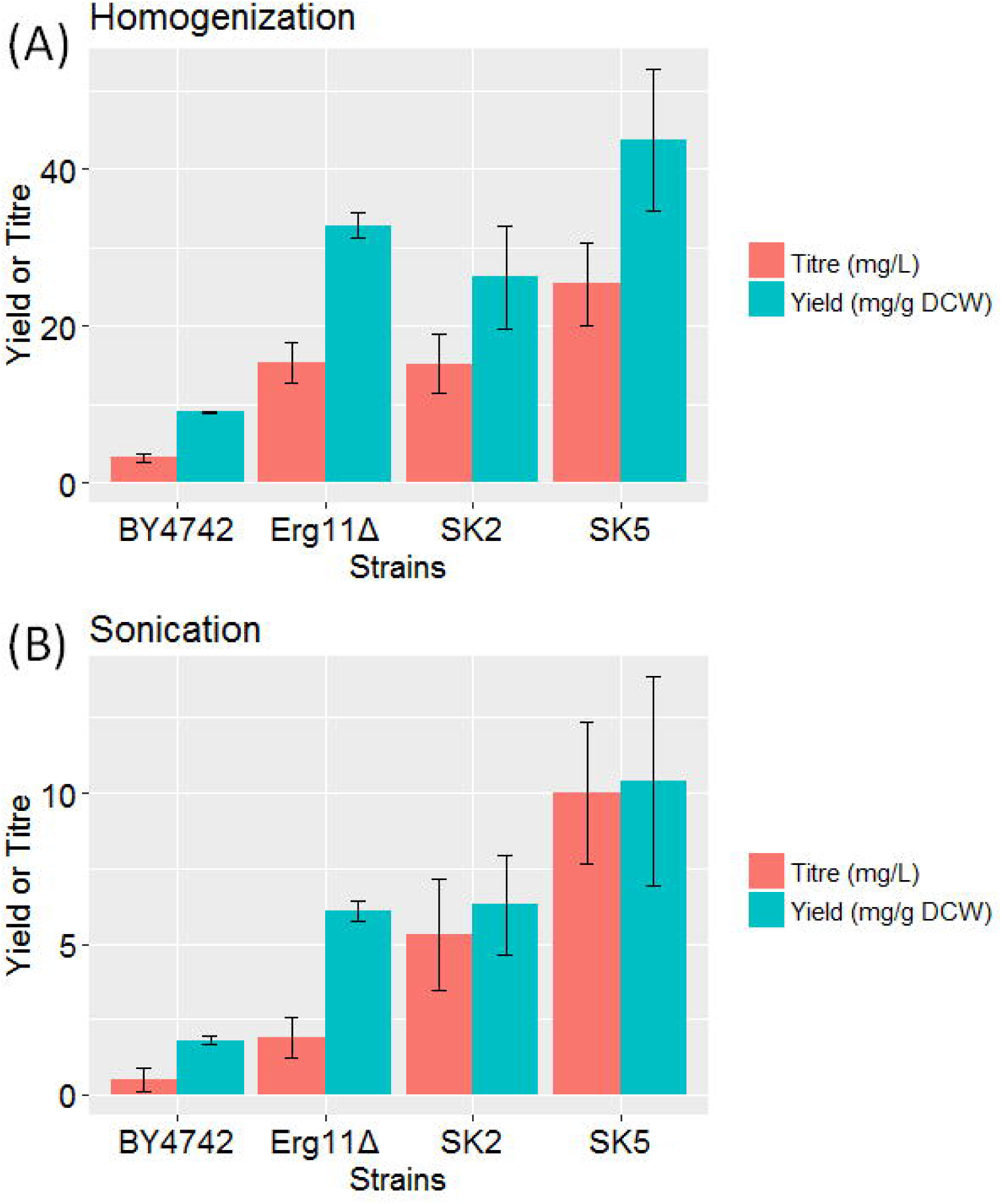
Culture condition optimization for squalene synthesis in yeast (A). Temperature optimization. (B). pH optimization.

### Effect of *ERG6* and *ERG11* gene deletion in squalene accumulation

Mutants of two enzymes from the downstream pathway has been chosen for screening of squalene accumulation and it has been observed that Erg11p diploid mutant has shown higher level of squalene than the Erg6p haploid mutant. Two wild-type strains BY4742 and CEN.PK2-1C has been chosen to compare and analyse squalene synthesis as CEN.PK2-1C has been a well-known strain for terpene synthesis with higher flux moving towards mevalonate pathway (Otero et al. 2010). As reported in several studies, CEN.PK2-1C has shown a better squalene yield which is 1.6-fold higher than BY4742. *ERG11* deletion strain has accumulated the highest amount of squalene of 6.1 mg/g DCW which is 3.4– fold, 3–fold and 1.9–fold higher than BY4742, CEN.PK2-1C and *ERG6* while *ERG6* deletion strain has shown 1.8–fold higher squalene yield than the parent control strain (Figure 4A).

**Figure 4.**
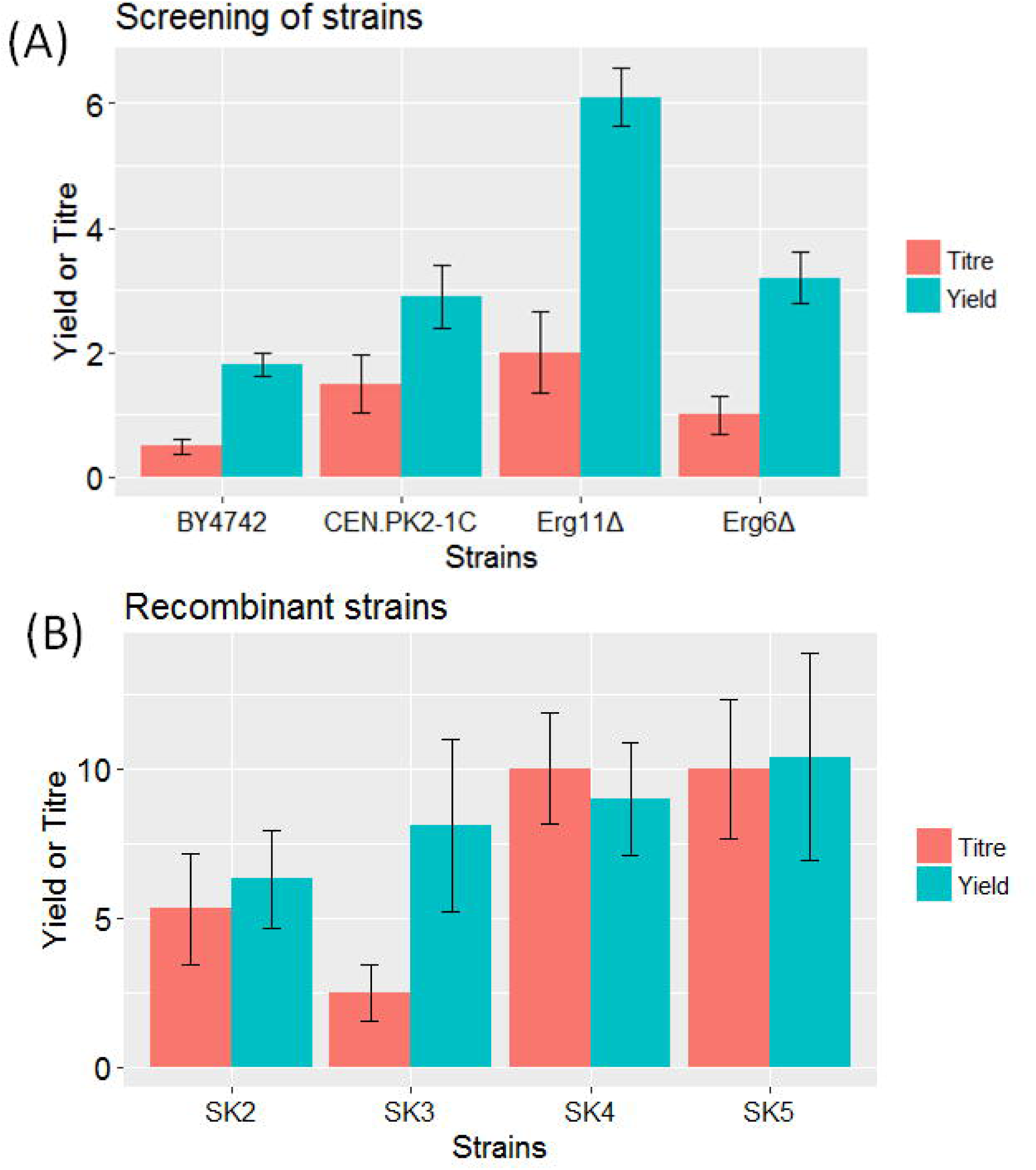
(A). Screening of different wild-type and mutant strains for squalene production. (B). Evaluation of extraction method in recombinant strain.

### Effect of over-expression of HMG-CoA reductase in squalene synthesis and its estimation

Further, to improve squalene synthesis, truncated HMG-CoA reductase has been overexpressed under the strongest inducible galactose promoter in two wild-type strains, BY4742 and CEN.PK2-1Cand the two mutant strains *ERG6Δ* and *ERG11Δ*. Overexpression of *tHMG1* has enhanced squalene accumulation more than five folds in all the strains. The combination of *ERG6Δ* and *ERG11Δ* with the *tHMG1* overexpression has also substantially improved squalene synthesis when compared with the wild-type strains. Squalene yield of 9 mg/g DCW and 10 mg/g DCW has been achieved in SK4 and SK5 strains respectively (Figure 4B) (Table S1).

Further, When the squalene extraction was carried out using homogenization method of cell lysis for the squalene hyper-producer SK5, we have achieved an yield of 43 mg/g DCW which is 5-fold higher over the parent control strain BY4742 which has given an yield of 9 mg/g DCW with homogenization (Figure 5B) (Table S2). *ERG6* has also been previously targeted for squalene improvement where Hmg2p overexpression combined with Erg6p deletion has not shown significant improvement in squalene content (Mantzouridou and Tsimidou 2010). Erg11p is a potential novel target enzyme chosen in this study to enhance squalene synthesis and it has been found to be a promising target for strain engineering towards terpene improvement.

**Figure 5.**
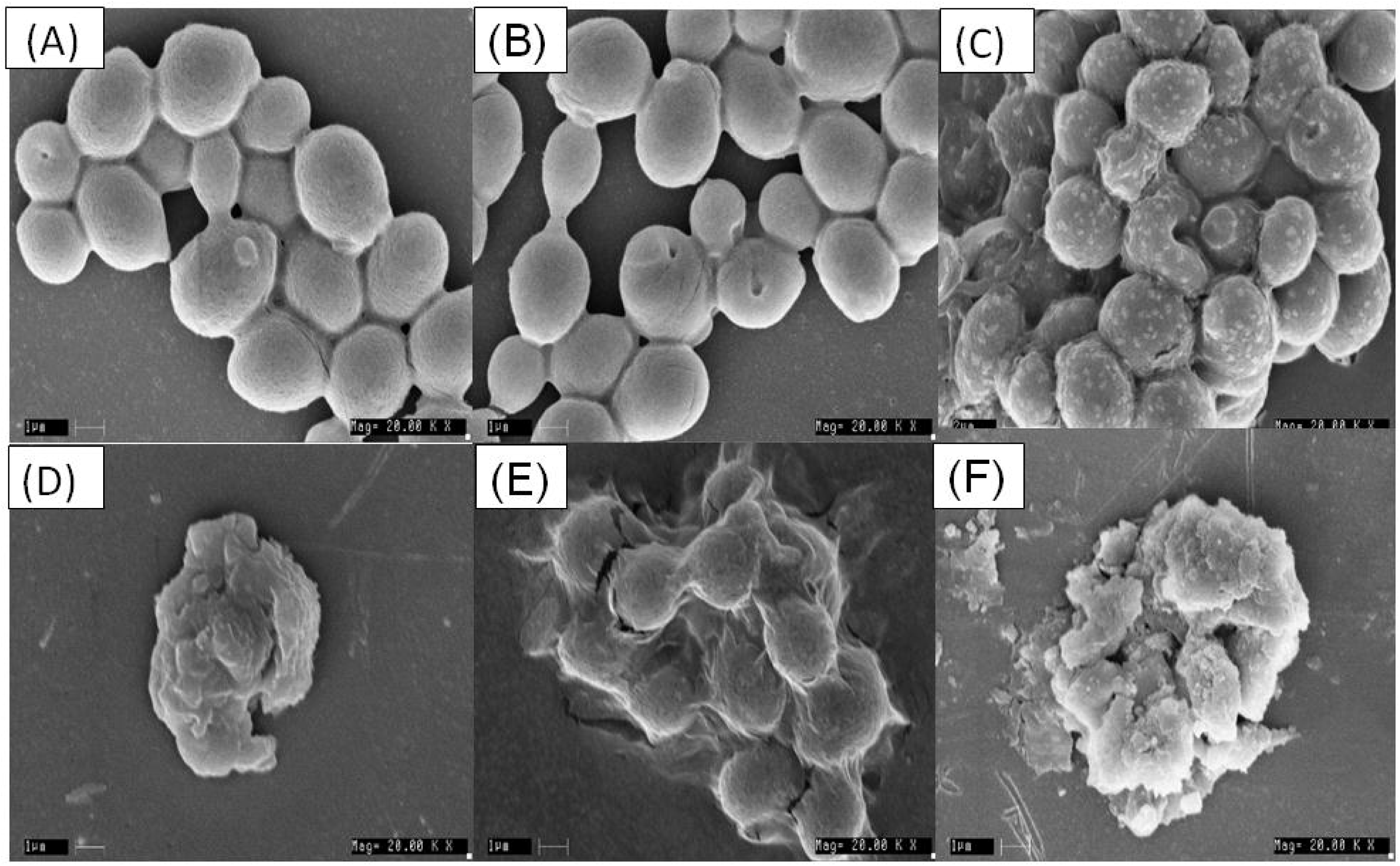
(A). Effect of overexpression of *tHMG1* gene combined with wildtype, *ERG6* knockout and *ERG11* knock-down background in squalene yield using sonication method of cell lysis. (B). Effect of overexpression of *tHMG1* gene combined with wildtype and *ERG11* knock-down background in squalene yield using homogenization method of cell lysis

The intensity of squalene band was found to be higher in SK5 in which *ERG11*Δ is combined with *tHMGl* over-expression when compared with SK2 and SK4. Meanwhile, ergosterol was detected in SK2 as well as in SK4 while it was found to be undetectable in SK5. *ERG11*Δ has reduced the ergosterol biosynthesis while *ERG6*Δ has shown lower ergosterol content (Figure S1). However, *ERG11*Δ strain might also have produced ergosterol which is not detected in the screening. Squalene was detected at higher levels in transformant strains when the carbon source was altered. The yeast cells provided with 1.8% galactose + 0.2% glucose has shown higher squalene yield when compared with the cells provided with 2% galactose as carbon source. Glucose is the most preferable carbon source in *S. cerevisiae* and the biomass is higher with glucose as a carbon source than galactose. Hence, when the cells were supplied with both 1.8% galactose and 0.2% glucose, it improved the biomass and also leading to squalene enhancement (Figure S1).

The cell dry weight was plotted against residual glucose concentration and the absorbance of cells at 600 nm. Cell dry weight was found to be 1.4 times lower in transformant wild-type strain than the wild-type strain while the cell dry weight is 1.75 times lower than the *ERG*6Δ and *ERG*11Δ mutant strains. This observation implies that the growth is compromised in the mutant strains due to the impaired ergosterol biosynthesis as the ergosterol is an integral component of cell membranes. Further, the biomass is reduced in the transformant strains due to the presence of the extra-chromosomal plasmid present inside the cells. The wild-type cells has shown a cell dry mass of 70 mg/g DCW at the end of 30 hours while the transformant cells has shown a cell dry weight of 50 mg/g DCW at the end of 30 hours. The cells were cultivated with an initial glucose concentration of 2% in synthetically defined medium. Residual glucose concentration has reduced to 0.2 % at the end of 30 hours whereas the glucose assimilation was found to be lower in case of transformant strain. The glucose concentration has reached 0.7 % at the end of 12 hours in the wild-type and mutant strains while it has reached 1 % in case of the transformant strain. In the mid time-course, glucose consumption has not been directed towards biomass production suggesting that the glucose has been used for energy maintanence rather than in the macromolecule synthesis. Transformant cells has been grown in glucose to analyze the biomass and cell dry weight. However, for enhancement of squalene the cells were grown in galactose based media (Figure S2).

### Effect of external squalene on growth of *S. cerevisiae*

Squalene has been included along with other media components at various concentration ranging from1.0-25.0 g/L. interestingly, squalene enhanced the growth of yeast cells in the selected range than control where there was no squalene addition. As the biomass has been increased up to 3.5-fold in the strains supplied with the squalene and the biomass increased with the increase in the squalene content in the medium (Figure S3), there might be a possibility that *S. cerevisiae* has used squalene as a carbon source. Squalene is not found to be lipotoxic to the normal wild-type cells even at higher concentration up to 25 g/L. Hence it could be produced in large amounts in the improved strains of *S. cerevisiae* while rendering the viability of the cells.

## Discussion

Sonication has been proven to be an efficient method for lipid extraction from several algal species such as *Chlorella sp*., *Nostoc sp*., and *Tolypothrix sp*. (Prabakaran and Ravindran 2011). Sonication causes cell-wall disruption by cavitation effect inside cells which in turn causes rupturing of cell walls by shock wave induction. The cell shearing is caused by high-frequency sound waves which leads to cavitation, a process involving the formation and collapse of gas bubbles which in turn leads to the conversion of sonic energy into mechanical energy as the gas bubble collapse increases the pressure and causes cell disruption. (Gogate 2011; Liu et al. 2013). Homogenization which is used for emulsification can also be used for efficient microbial cell wall disruption. During homogenization, the cells are forced through a valve and the high impact due to their interaction causes the disruption of cells (Clarke et al. 2010). Squalene has been conventionally extracted by chemical method of saponification that involves different percentage of KOH, pyrogallol and methanol. Saponification method of extraction has been developed for steryl esters that involves steps such as (i) addition of pyrogallol which acts as an antioxidant, (ii) heat treatment for cell lysis and (iii) methanol addition that prevents derivative formation (Adams and Parks 1968). However, the same method has also been used to extract non-saponifiable lipid fraction such as squalene which is present in cytosol and also sequestered in lipid particles and endoplasmic reticulum leading to poor yield of the compound when compared with squalene recovery by mechanical disruption of cells. The proposed methods in the current study combines lyophilization and sonication/homogenization of yeast cells. Lyophilization of cells has been shown to increase the squalene yield to 40-folds from *Torulaspora delbrueckii* species (Bhattacharjee and Singhal 2003). Lyophilization (freeze drying) technique has been employed in sample preparation as it makes the cells brittle thereby helping in efficient cell lysis by sonication (Dunstan et al. 1993). Hence, in our study the cells were harvested and subjected to lyophilization thereby aiding in the downstream extraction process.

Pyrogallol-Methanol-KOH extraction method that has been widely used for squalene and other terpene/terpenoid extraction from *S. cerevisiae* seems was found to be sub-optimal than mechanical cell lysis methods. The wild-type strain EGY48 has given a squalene yield of 0.6mg/g DCW using saponification with 60% KOH at 45°C (Mantzouridou and Tsimidou 2010). The parent strain YUG37 has shown a squalene yield of 0.16 mg/g DCW with acidic saponification at 70°C (Hull et al. 2014). The squalene yield in the wild-type strain is not reported in Shin et al. (Shin et al. 2012) while the squalene content of the wild-type strain used by Tokuhiro et al. (Tokuhiro et al. 2009) is not determined by the glass beads method whereas in the current study, we could extract and detect squalene in negligible amounts by applying both the methods using BY4742 as a parent strain.

One of the most common lipid extraction solvent system used is chloroform: methanol: water (1:2:0.8) (Dunstan et al. 1993) as squalene being a non-polar neutral lipid, we hypothesized chloroform-methanol solvent system will give higher squalene yields. The strong solvent system, chloroform and methanol combination has been chosen as it has increased lipid recovery over other solvent systems from several organisms including humans, microalgae as well as *S. cerevisiae* (Ferraz et al. 2004; Park et al. 2015). Chloroform being a lesser polar solvent and methanol being a polar solvent enhances the damage of cell wall thereby aiding in the release of squalene from the cells (Park et al. 2015). Chloroform-methanol solvent of 2:1 ratio has shown better squalene recovery when compared with other solvents such as petroleum ether and chloroform (Bhattacharjee et al. 2001). The same solvent system has also been used for squalene extraction from euglena (Kawaura et al. 1995). A 2:1 mixture of chloroform-methanol extracts the lipids more efficiently over the other solvent systems (Christie 1982). The cells were dispersed in the chloroform-methanol solvent system prior to sonication and was subjected to overnight extraction in shake flasks after sonication to further increase the shearing of cells.

Squalene synthesis was found to be drastically reduced in the cells grown at 40°C due to significantly lower cell yields. The optimum pH and temperature has shown slight variations from the optimal growth conditions for the yeast cells probably due to the reason that the enzymes has different favorable conditions and the ones involved in squalene biosynthesis might be having a slightly different optimal requirements for its efficient activity.

Two enzyme mutant strains harbouring *ERG6*, non-essential gene deletion and *ERG11*, essential gene deletion, respectively, in the downstream sterol biosynthesis pathway has been selected to analyze the squalene synthesis under ergosterol biosynthesis impairment. The primary genes in the heme-dependent pathway are essential while the genes following *ERG6* are nonessential indicating that the sterol pathway intermediates such as fecosterol, episterol and zymosterol might take up the role of ergosterol in plasma membrane in its absence (Alvarez-vasquez et al. 2011). Erg11p has been selected as a mutant target as all the enzyme deletion at or above the Erg11p renders the cells auxotrophic for ergosterol and Erg11p is the terminal key essential enzyme in the pathway while Erg6p is the foremost non-essential enzyme in the pathway as the gene knock-out strains of the enzymes below Erg6p are viable (Sturley 2000). Further, Erg11p requires three molecules of NADPH for its activity and *ERG11* knock-down in turn can reduce the activity of Erg11p and improve the availability of NADPH for HMG-CoA reductase and squalene synthase activity. As *ERG11* being an essential gene, its knockout has constrained the flux towards ergosterol pathway thereby leading to squalene accumulation while *ERG6* being a non-essential gene hasn’t shown a notable improvement in squalene than *ERG11* mutant (Figure 4B) (Table S1). In Figure S1, *ERG11*Δ has shown negligible ergosterol formation and this might be due to the reason that the minimal amount of ergosterol synthesized in *ERG11*Δ strain would have formed complexes with sphingolipids and get embedded in the membrane which is hard to break open than the steryl-esters in lipid droplets.

Initial efforts have been made to improve squalene by overexpression of HMG-CoA reductase enzyme in yeast cells. Hmg1p is an integral membrane protein in the endoplasmic reticulum with two domains, viz a transmembrane domain and a catalytic domain. The expression of the whole protein Hmg1p has lead to karmellae formation while the increased expression of the catalytic domain of Hmg1p exclusively has shown to channelize the carbon flux towards mevalonate pathway and the squalene accumulation in strains (Wright et al. 1988; Polakowski et al. 1998).

Components with molecular weight (Mr) more than 760 has been shown to penetrate through the cells. *S. cerevesiae* cell wall has a wide range of porosity at exponential-growth phase that is permeable for molecules with size upto 70 kDa (Klis et al. 2014). Squalene with the molecular weight of 410 daltons can readily move through the membrane. Squalene can also be used as a carbon source by several microorganisms such as eubacteria (*Corynebacterium, Pseudomonas,* or *Arthrobacter*) (Ghimire et al. 2016). Although squalene is lipotoxic to cells defective in lipid droplet formation, it is not growth-inhibitory in normal wild-type cells. This could be due to the fact that the excess squalene are sequestered and stored in lipid droplets along with triacyl glycerides and steryl-esters (Spanova et al. 2012) (Figure 6).

**Figure 6.**
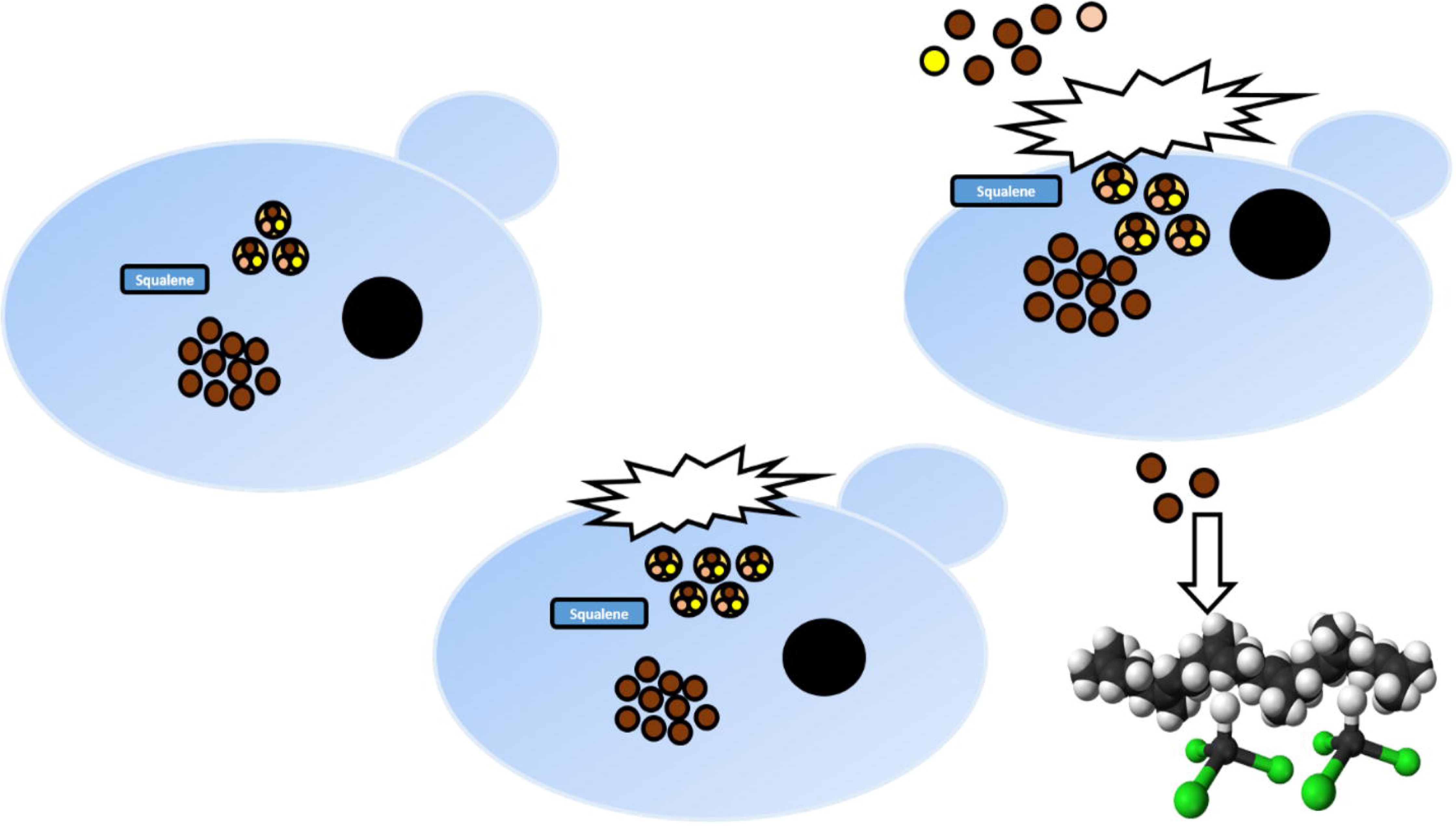
Squalene release from lipid droplets

To conclude, sonication and homogenization has been used as a cell-disruption method for efficient extraction of squalene from *S. cerevisiae* strains. Both qualitative as well as quantitative methodologies have been established to observe squalene synthesis in *S. cerevisiae.* The mutant strains in the downstream ergosterol pathway and *tHMG1* overexpressed strains were also used to validate the robustness of squalene analysis. Homogenization based mechanical disruption was observed to yield higher squalene and the SEM analysis corroborates these findings. The synergistic effect of *ERG11* downregulation and *tHMG1* over-expression has lead to significant increase in squalene yield. The proposed homogenization method could also be exploited for the analysis of other terpenes’ biosynthesis using *S. cerevisiae.* It was further proved that when the galactose induction was coupled to glucose addition there was a significant improvement in the squalene synthesis in the tHMG1 overexpressed strain under galactose promoter. Also it has been observed the *S. cerevisiae* have not only tolerated 25 g/L of external squalene but also improved the biomass in the presence of squalene.

## Acknowledgements

The acknowledgements come at the end of an article after the conclusions and before the notes and references. KP acknowledges the Department of Biotechnology, India, for the award of research fellowship. SM wish to acknowledge the financial support provided by Science Engineering and Research Board (SERB), India. We acknowledge Prof. Ram Rajasekharan (Lipid Science Dept., CSIR-CFTRI) for providing us the strains and the plasmid, pYES2/NTC. The authors are grateful to Mr. K. Anbalagan, CIFS, CFTRI for his help in SEM analysis and Mr. P. Mukund Lakman, CIFS, CFTRI for his help during HPLC.

## Supporting information

Additional Supporting Information may be found in the online version of this article:

### Figure S1

Qualitative TLC screening for squalene and ergosterol in the mutant and transformant

### Figure S2

Growth behavior pattern of A. Wildtype B. Transformant strain, SK2 C. *ERG*6Δ D. *ERG*11Δ in synthetically defined media.

### Figure S3

Effect of squalene at various concentrations on growth of *S. cerevisiae*.

### Table. S1

Squalene yield and fold improvement from different strains using sonication

### Table. S2

Yield and fold improvement from squalene over-producer using homogenization Tables

